# The positive evidence bias in perceptual confidence is not post-decisional

**DOI:** 10.1101/2020.03.15.991513

**Authors:** Jason Samaha, Rachel Denison

**Affiliations:** University of California, Santa Cruz; New York University

## Abstract

Confidence in a perceptual decision is a subjective estimate of the accuracy of one’s choice. As such, confidence is thought to be an important computation for a variety of cognitive and perceptual processes, and it features heavily in theorizing about conscious access to perceptual states. Recent experiments have revealed a “positive evidence bias” (PEB) in the computations underlying confidence reports. A PEB occurs when confidence, unlike objective choice, over-weights the evidence for the chosen option, relative to evidence against the chosen option. Accordingly, in a perceptual task, appropriate stimulus conditions can be arranged that produce selective changes in confidence reports but no changes in accuracy. Although the PEB is generally assumed to reflect the observer’s perceptual and/or decision processes, post-decisional accounts have not been ruled out. We therefore asked whether the PEB persisted under novel conditions that eliminated two possible post-decisional accounts: 1) post-decision evidence accumulation that contributes to a confidence report solicited after the perceptual choice, and 2) a memory bias that emerges in the delay between the stimulus offset and the confidence report. We found that even when the stimulus remained on the screen until observers responded, and when observers reported their choice and confidence simultaneously, the PEB still emerged. Signal detection-based modeling also showed that the PEB was not associated with changes to metacognitive efficiency, but rather to confidence criteria. We conclude that once-plausible post-decisional accounts of the PEB do not explain the bias, bolstering the idea that it is perceptual or decisional in nature.

## Introduction

Human perceptual decision making not only results in a choice about what is perceived but can also produce a sense of confidence in the accuracy of that choice. Several lines of inquiry suggest that the same source of evidence informs both the choice and one’s confidence in that choice. For instance, subjective reports of confidence are well-correlated with choice accuracy across a variety of perceptual and mnemonic tasks (Ais et al., 2016; Grimaldi et al., 2015; Kiani et al., 2014; Samaha & Postle, 2017; Song et al., 2011). Moreover, certain neurons in monkey parietal cortex encode both the choice and confidence level of the animal (Kiani & Shadlen, 2009). The close relation between choice accuracy and confidence has been formalized in computational approaches that define confidence as the probability of being correct, thereby casting confidence as an optimal readout of choice uncertainty (Kepecs et al., 2008; Kiani et al., 2014; Meyniel et al., 2015; Sanders et al., 2016).

In contrast, a number of recent experiments have demonstrated a bias in confidence reports that renders confidence dissociable from choice accuracy. The so-called “positive evidence bias” (PEB) refers to the finding that confidence seems to over-weight the evidence in favor of the chosen option (the “positive evidence”), whereas the difficulty of the choice itself is governed by the balance of evidence between choice alternatives (Koizumi et al., 2015; Odegaard et al., 2018; Peters et al., 2017; Rausch et al., 2017; Samaha et al., 2016, 2017, 2019; Zylberberg et al., 2012). For example, Koizumi et al. (2015) demonstrated, in a motion-direction discrimination task, that increasing the number of dots moving in the direction of the correct choice while simultaneously increasing the number of dots moving randomly does not change accuracy but increases confidence.

We have recently demonstrated the PEB in orientation discrimination and estimation tasks. In our protocol, a single luminance-modulated grating is embedded in white noise, and we examine the effect of increasing both the noise and the grating contrast. We observed that proportional increases in grating and noise contrast selectively boosts confidence ratings, but not discrimination or estimation accuracy (Samaha et al., 2016, 2019). Given the importance of accurate confidence for guiding appropriate behaviors (Desender et al., 2018; Folke et al., 2017), and the role of confidence effects in theoretical accounts of conscious perception (Brown et al., 2019; Dehaene & Changeux, 2011; Lau & Rosenthal, 2011), an understanding of why confidence diverges from accuracy in the PEB is warranted.

Theoretically, the PEB in orientation tasks could emerge at perceptual stages of processing if the orientation strength appears stronger or more visible to the observer, thereby causing the change in confidence. The PEB could occur at decision stages, for example, if stimulus information other than the orientation percept is used to inform confidence. A third option is that the bias emerges after the initial perceptual decision is made. Here, we consider two such post-decisional accounts of how confidence and accuracy can dissociate and test whether they explain the PEB.

The first account is termed post-decision evidence accumulation (Navajas et al., 2016). It is based on the idea that choice evidence continues to accumulate even after the initial decision is made. If, as is the case with all prior experiments on the PEB, confidence is solicited *after* the stimulus choice, then confidence can be based on different levels of evidence than the choice, leading to dissociations. This account has been leveraged to explain changes of mind, among other confidence-related results (Navajas et al., 2016). The second account is that the PEB does not arise in perceptual decision-making, *per se,* but is a bias that arises in short-term memory. This is plausible because experiments demonstrating the PEB so far have used relatively short, fixed-duration stimulus presentations, followed by choice and confidence responses. Thus, the confidence judgment is typically rendered half a second or more after the stimulus was perceived.

We test both of these accounts in a new experiment in which observers issue their choice and confidence reports at the same moment in time with a single key press (eliminating post-decisional evidence accumulation) and where confidence is reported while the stimulus is still visible (eliminating any memory-based bias). Using a signal detection theory (SDT) analysis, we find that the PEB persists under these conditions and that the bias is linked to confidence criteria, rather than metacognitive efficiency. These results suggest that the PEB arises during perception or decision-making, rather than during a post-decisional stage.

## Materials and Methods

### Participants

26 participants (age range: 18-35 years; 17 female) from the University of Wisconsin-Madison community completed the experiment. 23 of the participants provided data deemed suitable for hypothesis testing (see *Staircase Procedure*). All participants reported normal or corrected visual acuity, provided written informed consent, and were compensated monetarily. Sample size was based on prior experiments we have done looking at the PEB across two or more conditions (Samaha et al., 2016, 2019). This experiment was conducted in accordance with the University of Wisconsin Institutional Review Board and the Declaration of Helsinki.

### Data Availability

In accordance with the practices of open science and reproducibility, all task scripts, raw data, and code used in the present analyses are freely available through the Open Science Framework (https://osf.io/5m9wd/)

### Stimuli

Visual stimuli were composed of a sinusoidal luminance grating (1.1 cycles per degree, zero phase) embedded in white noise and presented centrally within a circular aperture (2.5 degrees of visual angle [DVA]). The orientation of the grating was randomly chosen on each trial to be 45° or −45° tilted from vertical. The noise component of the stimulus was created anew on each trial by randomly sampling each pixel’s luminance from a uniform distribution. A fixation point (light gray, 0.19 DVA) was centered on the screen throughout the trial. Stimuli were presented on a gray background on an iMac computer screen (52 cm wide by 32.5 cm tall; 1920 by 1200 pixel resolution; 60 Hz refresh rate) using PsychToolbox 3 (Kleiner et al., 2007; Pelli, 1997) running in MATLAB 2015b (MathWorks, Natick, MA) viewed from a chin rest at a distance of 62 cm.

### Staircase Procedure

The PEB can be demonstrated by embedding gratings in noise under two different conditions (Fig. 1A): one high contrast grating averaged with high contrast noise (which we will refer to as *high positive evidence* or high PE) and one low contrast grating averaged with low contrast noise (*low positive evidence* or low PE; Samaha et al., 2016, 2019). If there is a PEB, then the high PE stimulus will produce higher confidence ratings, but not higher accuracy. We therefore started each main task with a staircase procedure designed to find two levels of grating contrast that produce equal levels of discrimination accuracy when embedded in low and in high contrast noise. Specifically, we started with a 50% and a 100% Michelson contrast noise patch and used the Quest procedure as implemented in PsychToolbox 3 to find a grating contrast that produced ~75% accuracy when averaged with the 50% noise patch and another contrast level that produced the same accuracy when averaged with the 100% noise patch. In previous work, we had used a single staircase to find a grating contrast threshold that, when averaged with 100% noise should produce ~75% accuracy and then simply halved the contrast of the grating and the noise to make the low PE stimulus. Although this procedure previously worked to match accuracy at the group level (as predicted by Weber’s law), here we matched accuracy empirically at the individual participant level by separately staircasing high and low PE stimuli. To this end, we ran 20 practice trials followed by 200 trials of the staircase before each of the two main task conditions (fixed duration vs. response-dependent; see *Main Task Procedure*). High PE and low PE staircases were interleaved and the final grating contrast for the low and high PE stimuli was computed as the mean of the Quest posterior distribution. The thresholds from some participants, however, deviated substantially from the predicted value of 50% lower grating contrast in the low PE condition. To ensure that variability in the efficacy of the staircase did not overly influence the results, we excluded three subjects whose low PE thresholds were less than 20% or greater than 80% of their high PE thresholds. The staircase followed the same task structure as the main tasks (described next), with the only exception that the contrast of the grating component of the stimulus was adapted using Quest.

**Figure 1.**
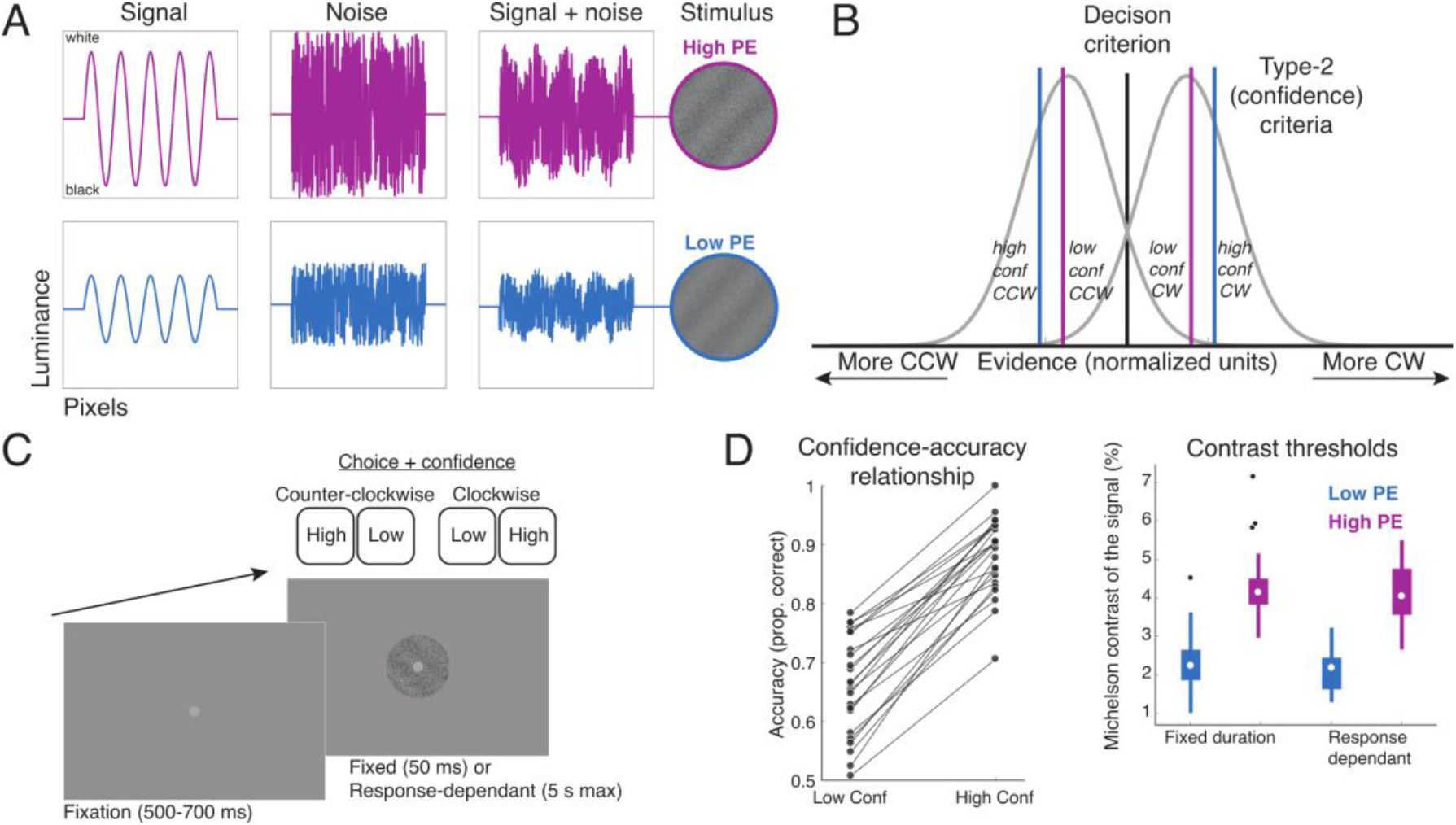
**A.** Schematic of stimuli used in the positive evidence (PE) manipulation. High PE stimuli were composed of 100% contrast white noise averaged with a higher contrast grating (contrast thresholds in panel D, right), whereas low PE stimuli contained 50% contrast noise averaged with a lower contrast grating. **B.** SDT model of confidence and performance. Gaussian distributions represent internal evidence for clockwise (CW) and counter-clockwise (CCW) stimuli across trials. Type-2 (confidence) criteria (colored vertical lines) are exceeded when a certain level of evidence (distance from decision criterion) is passed. The PEB could arise if, on high PE trials, the confidence criteria shift closer to the decision boundary, leading to more frequent reports of “high confidence” without changes in accuracy. Note that such a shift would occur in the normalized evidence space if, for example, the means and variances of the absolute evidence distributions scaled for high PE stimuli but the absolute confidence criteria did not change (not shown). **C.** Task schematic. On each trial, a high or low PE stimulus tilted either 45 or −45 degrees from vertical was presented. In different blocks, the stimulus was either presented for a fixed duration of 50 ms, or, until a response was made (up to 5 seconds). Choice (CW or CCW) and confidence level (high or low) was given with a single button press. **D.** Left, accuracy is higher for decisions endorsed with high confidence. Right, boxplots of contrast thresholds for each PE level and task, determined from a pre-task adaptive procedure. Note that these are the contrast levels of the gratings prior to averaging with 100% (high PE) or 50% (low PE) contrast white noise.

### Main Task Procedure

To test whether the PEB persisted when the target stimulus remained onscreen until the participant’s response, we tested both a fixed duration condition in which the stimulus duration was 50 ms and a response-dependent condition in which the stimulus was displayed until a response was made. The only difference between conditions was the duration of the target stimulus.

Each trial began with an inter-trial interval (ITI) randomly drawn from a uniform distribution of durations between 500 and 700 ms. During the ITI, only the central fixation point was displayed. A target stimulus followed, which was randomly selected on each trial to be 45° or −45° from vertical and have high or low PE. Choice (clockwise/counter-clockwise) and confidence (two levels: high or low) were indicated with a single keypress. With their left hand, participants used the “F” and “D” key to indicate counter-clockwise tilt with low and high confidence respectively. A right-hand response using the “J” or “K” key indicated clockwise tilt, low and high confidence respectively. In this way, the confidence report was made with the same amount of accumulated evidence as the orientation choice (Fig. 1C). Responses were required within 5 seconds of stimulus onset and trials with late responses were repeated at the end of the block. Participants completed 3 blocks of 80 trials each of the fixed duration condition, and then 3 blocks of 80 trials each of the response-deponent condition, or vice versa (counterbalanced across participants). Before starting either duration condition for the first time, the staircase was run to find high and low PE thresholds for that condition. Those threshold contrast values were then used for the main task.

### Model-free analysis

For each combination of PE and stimulus duration, we computed mean confidence ratings and accuracy (proportion correct) across trials. We submitted these measures to separate 2-by-2 repeated-measures ANOVA with PE (high or low) and stimulus duration (fixed or response-dependent) as factors. If the PEB emerges in memory, then we should see a confidence bias in the fixed duration stimulus condition, but not in the response-dependent condition, corresponding to an interaction between PE and duration when predicting confidence. If the PEB is present in both conditions, we expect a main effect of PE on confidence. Additionally, we expect no main effect of PE on the proportion of correct discrimination responses if we adequately controlled performance. Lastly, we analyzed the mean response time (RT) in all conditions to determine whether changes in confidence (without changes in accuracy) was associated with decision duration, as predicted by some drift-diffusion models of confidence (Kiani et al., 2014; Kiani & Shadlen, 2009; Zylberberg et al., 2016). Using median RT did not produce any different results.

### Model-based analysis

In addition to a model-free analysis (which ensures that results are not based on any particular model assumptions), we also used a SDT modeling framework (Fig. 1B) to assess the impact of PE (high or low) and stimulus duration (fixed or response-dependent) on behavior. The model, called Meta-d’ (Fleming & Lau, 2014; Maniscalco & Lau, 2012), estimates type-1 sensitivity (d’), 2) type-2 criteria (confidence criteria), which reflect how much stimulus evidence is needed to commit to a high or low confidence response, and 3) metacognitive efficiency (meta-d’ – d’), which reflects how well confidence ratings distinguish between correct and incorrect responses relative to an ideal observer with no information loss between the type-1 (choice) and type-2 (confidence) decisions. Meta-d – d takes on negative values when metacognitive sensitivity is worse than what would be expected by an ideal (according to SDT) metacognitive observer, whereas values close to zero indicate no deviation from optimality. We derived a single estimate of confidence criteria for each observer by averaging over the absolute values of the criteria associated with clockwise and counter-clockwise stimuli. The resulting metric indicates normalized (z) evidence criteria needed to commit to a “high confidence” decisions; lower values therefor indicate more liberal type-2 criteria (less evidence is needed). The model was estimated from choice and confidence ratings for each participant using a Bayesian model called HMeta-d (Fleming, 2017) wherein the calculation of d’ follows the model of (Lee, 2008). This open-source software is available at (https://github.com/metacoglab/HMeta-d). We used the MATLAB implementation (function *fit_meta_d_mcmc.m*), which implements a Markov-Chain Monte Carlo sampling procedure to estimate posterior distributions over parameters. We ran the function with 3 chains, 1,000 burn-in samples, and 10,000 recorded samples in each chain.

We then entered each of the three parameters (confidence criteria, meta-d’ – d’, and d’) into separate 2-by-2 repeated-measures ANOVA with PE (high or low) and stimulus duration (fixed or response-dependent) as factors. If increasing PE boosts confidence ratings in both stimulus duration conditions, we expect a main effect of PE on confidence criteria and possibly on meta-d’ – d’ (if the PEB also alters metacognitive efficiency). Specifically, higher confidence ratings would correspond to lower confidence criteria, i.e. less evidence required to make a “high confidence” response. If, on the other hand, the PEB is caused by a memory bias, then confidence measures should interact with stimulus duration, as PEB would only occur in the fixed duration condition where judgments are made on the basis of a memory representation. If our staircase procedure adequately controlled for accuracy, we expect no effects of PE on d’.

## Results

### Staircase thresholds

Contrast thresholds from the staircase procedure conformed well to Weber’s law: in the fixed duration staircase, mean (±SEM) Michelson contrast for the low PE (2.33 ± 0.15%) and high PE (4.35 ± 0.20%) produced a ratio of 53 ± 2.5%. The response-dependent staircase produced threshold estimates for low PE (2.12 ± 0.12%) and high PE (4.12 ± 0.16%) with a ratio of 51 ± 2.5%. Threshold contrasts were numerically, though not significantly (p=0.18), lower for the response-dependent task, indicating that slightly lower thresholds are achieved when the observer has unlimited time to view the stimulus (Fig. 1D).

### General confidence behavior

Before addressing our main hypotheses, we confirmed that each participant used their confidence responses appropriately in that higher confidence was associated with higher accuracy. Collapsing across PE levels and task, every participant had higher mean accuracy on trials endorsed with high, as compared to low, confidence (Fig. 1D), *t*(22) = 16.56, *p* = 6.5^−14^.

### Stimulus viewing time

Since viewing time was controlled by the observer in the response-dependent condition, average stimulus duration varied from person-to-person between a range of 0.7 (min) to 1.8 (max) seconds, with a mean (SD) across subjects of 1.05 (0.30) seconds. In the fixed-duration condition, stimuli were always presented for 50 ms.

### Model-free results

These data are shown in Fig. 2. We found no main effect of PE on discrimination accuracy (*F*(1,22) = 0.10, *p* = 0.756) nor an effect of stimulus duration (*F*(1,22) = 2.01, p=0.170), indicating that the staircasing procedure effectively equated discriminability across PE levels and tasks. Additionally, there was no interaction between PE and duration when predicting accuracy (*F*(1,22) = 0.05, *p* = 0.833). Consistent with the PEB, however, we observed a significant main effect of PE on mean confidence ratings (*F*(1,22) = 5.08, *p* = 0.034): high PE was associated with higher confidence (consistent with a more liberal threshold for reporting high confidence). This main effect argues against post-decision evidence accumulation as the source of the PEB. Importantly, PE and duration did not interact to predict confidence (*F*(1,22) = 0.4, *p* = 0.532), suggesting that the PEB persists even when no memory is required. We also observed a main effect of duration on confidence (*F*(1,22) = 10.95, *p*=0.003), with lower confidence when the duration was response-dependent.

**Figure 2.**
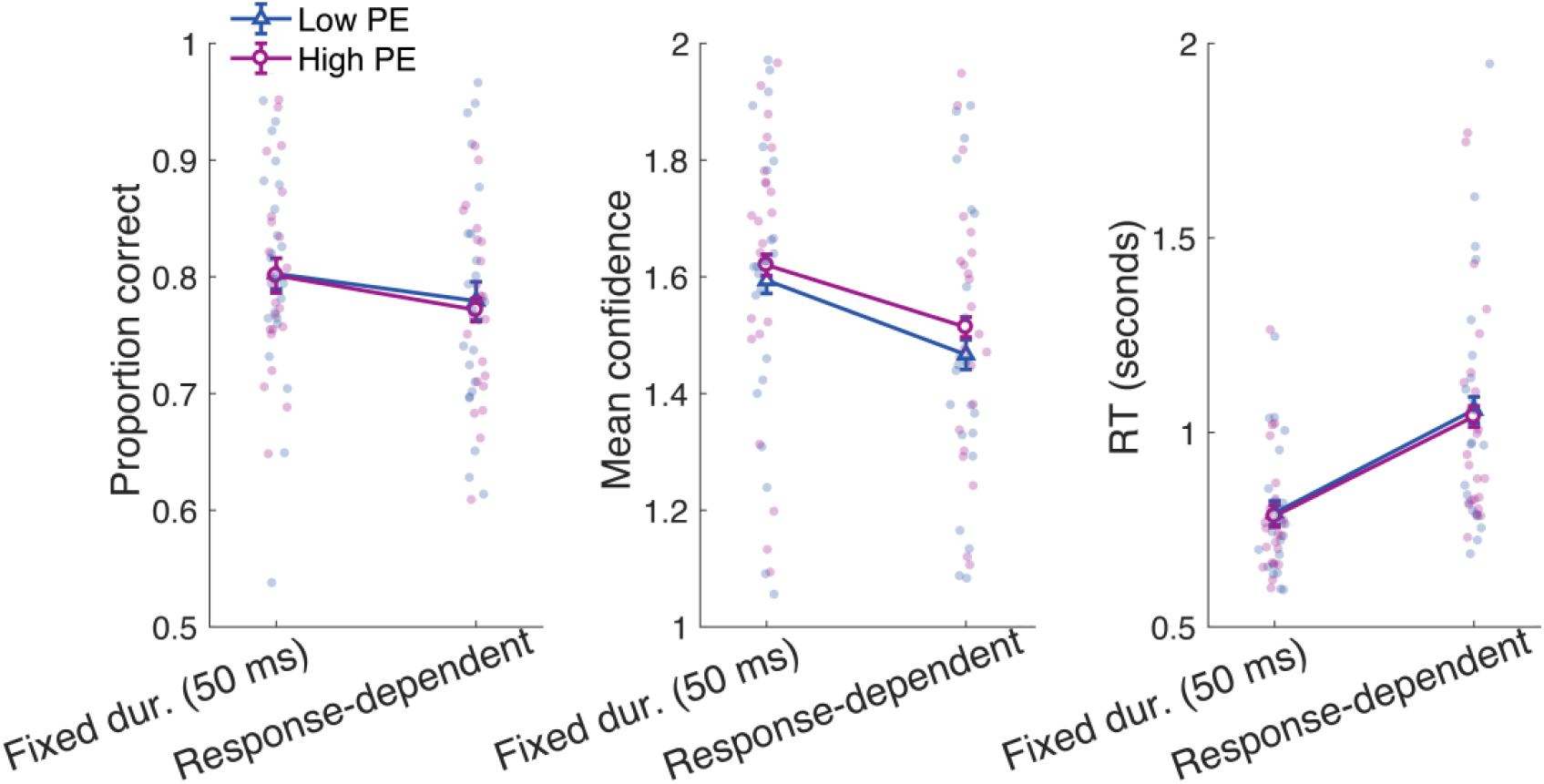
Model-free results depicting orientation discrimination accuracy (left), average confidence rating (middle), and response times (right) as a function of PE (high or low) and stimulus duration (fixed or response-dependent). A significant main effect of PE on confidence without an effect on accuracy confirms the PEB. A lack of interaction between PE and duration for confidence ratings suggests the bias is not task-dependent. No PE effect was observed on RT, though a duration effect is evident. Error bars show ±1 within-subject SEM (Morey, 2008); dots are individual observers.

To compliment the accuracy and confidence results, we also assessed RTs as a function of PE and stimulus duration (Fig. 2). There was no main effect of PE (*F*(1,22) = 1.72, *p* = 0.203), indicating that RT changes did not accompany PE-related changes in confidence. There was also no PE-by-duration interaction for RT (*F*(1,22) = 0.16, *p* = 0.696). However there was a clear main effect of duration (*F*(1,22) = 19.84, *p* = 0.0002), reflecting the fact that participants took longer to respond when the stimulus viewing duration was under their control.

### Model-based results

Using the SDT model of metacognition, Meta-d’, we estimated type-1 sensitivity (d’), type-2 criteria, which determine how much evidence is needed to endorse a response with high confidence, and type-2 sensitivity adjusted for type-1 sensitivity (Meta-d’ – d’). These data are summarized in Fig. 3. The staircase procedure effectively matched perceptual sensitivity across PE conditions. We found no main effect of PE on d’ (*F*(1,22) = 0.38, *p* = 0.541). There was also no main effect of stimulus duration on d’ (*F*(1,22) = 1.42, *p* = 0.245), indicating that the staircase matched performance between the fixed and response-dependent duration conditions. Duration and PE did not interact to predict d’ (F(1,22) = 0.07, *p* = 0.794).

**Figure 3.**
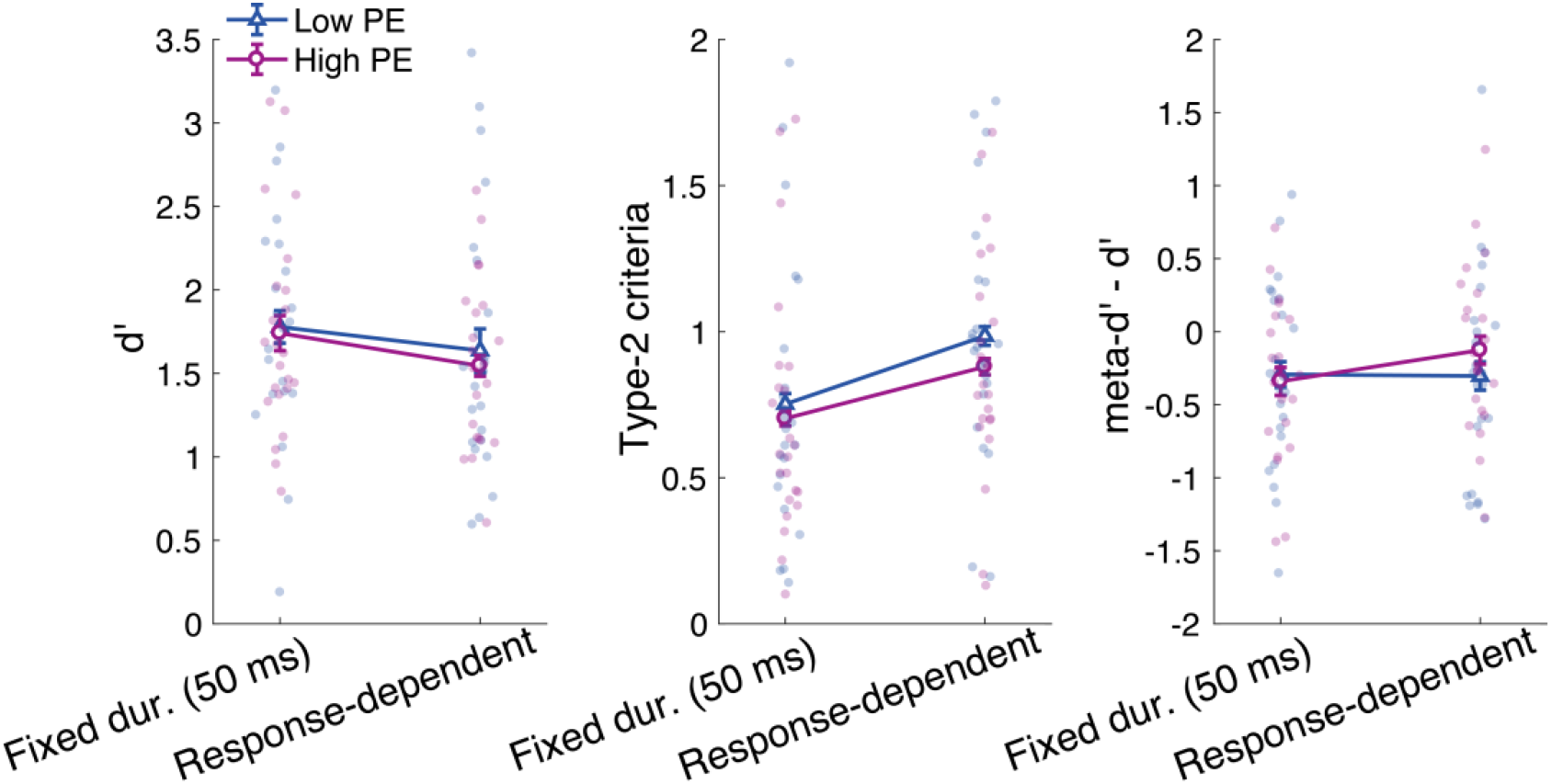
Model-based results showing d’ (left), type-2 criteria (middle), and metacognitive efficiency (meta-d’ – d’; right) as a function PE (high or low) and stimulus duration (fixed or response-dependent). A main effect of PE on type-2 criteria indicates a more liberal criteria with high PE and no effect on d’ confirms the PEB. A lack of interaction on type-2 criteria suggest the PEB is not task-dependent, and a lack of any effects on metacognitive sensitivity suggests the PEB is driven by a bias in overall confidence level rather than a change in the relation between confidence and accuracy. Error bars show ±1 within-subject SEM (Morey, 2008); dots are individual observers.

In contrast to the lack of effects on type-1 performance, the PE manipulation produced a significant main effect on type-2 criteria (*F*(1,22) = 10.38, *p* = 0.0039). High PE caused a more liberal type-2 criteria, indicating that less evidence was required to respond with high confidence. This main effect, coupled with a lack of significant interaction with stimulus duration (*F*(1,22) = 1.31, *p* = 0.265), suggests that the PEB does not depend on the stimulus being presented for a short duration. If anything, the type-2 bias was stronger in the response-dependent duration condition (Fig. 3). These findings indicate that the PEB does not arise in memory. In addition, we observed a significant main effect of stimulus duration on type-2 criteria, such that participants used more conservative type-2 criteria in the response-dependent duration task compared to the fixed duration task (*F*(1,22) = 16.52, *p* = 0.0005). To our knowledge, this is the first report of changes in type-2 criteria with stimulus duration.

The effects of PE on confidence reports were specific to type-2 criteria. For metacognitive efficiency (meta-d’ – ‘d), which describes how well confidence tracks performance while accounting for task difficulty, we observed no main effect of PE (*F*(1,22) = 0.32, *p* = 0.578) nor of task (*F*(1,22) = 0.78, *p* = 0.385), and there was no interaction (*F*(1,22) = 1.41, *p* = 0.248). This suggests that the PEB is purely an overall confidence bias and does not relate to the introspective accuracy of confidence judgments.

## Discussion

We investigated a previously documented bias in confidence ratings that is thought to occur because of an over-reliance of confidence computations on evidence for the chosen alternative (Koizumi et al., 2015; Maniscalco et al., 2016; Odegaard et al., 2018; Peters et al., 2017; D. Rahnev et al., 2011; D. A. Rahnev et al., 2012; Samaha et al., 2016, 2017, 2019; Zylberberg et al., 2012, 2014). We tested the PEB under novel circumstances where memory effects and post-decision evidence accumulation could be ruled out as explanations of confidence reports. Our main finding, evident in both a model-free analysis and a SDT-based model of confidence behavior, is that the PEB persisted under these new circumstances. We can therefore rule out memory and post-decision evidence accounts of the PEB. A further novel result is that the PEB is not associated with a reduction of metacognitive efficiency (the extent to which confidence ratings track decision accuracy, controlling for performance) but was accounted for by a shift in type-2 criteria, such that less evidence was needed to endorse a decision with high confidence when we increased both signal and noise in the stimulus. (Note that this type-2 criteria shift is measured in a variance-normalized evidence space and may or may not correspond to a change in the absolute amount of evidence needed to commit to a “high confidence” response).

Subjective reports of perception, such as confidence or visibility ratings, often figure as evidence in debates about the nature of conscious perception, the role of consciousness in behavior, and the neural correlates of consciousness (Block, 2011; Brown et al., 2019; de Lafuente & Romo, 2005; Dehaene & Changeux, 2011; Hesselmann et al., 2011; King & Dehaene, 2014; Lamme, 2003; Lau & Passingham, 2006; Samaha, 2015; Vandenbroucke et al., 2014). The phenomenon of blindsight has been particularly informative in the study of conscious perception precisely because it dissociates subjective and objective aspects of perception (Azzopardi & Cowey, 1997; Cowey & Stoerig, 1995; Overgaard, 2011; Weiskrantz, 1986). Whenever similar, albeit far less dramatic, dissociations occur in typical individuals, as is the case in the PEB, it raises the possibility of insight into the mechanisms involved in subjective conscious perception independent of objective task performance. However, such insight is only meaningful for perception if the change in the subjective report actually reflects something about an individual’s conscious experience, rather than a change in post-perceptual processes. By ruling out memory effects and post-decision evidence accumulation as sources of the PEB, our results indicate that a perceptual account of the PEB remains viable – along with the possibility that the bias is caused at some decisional, rather than perceptual stage of processing.

If the bias is genuinely perceptual, which future work will continue to address, it would suggest that different computations underlie subjective visual appearance and objective decision making. The PEB is thought to occur because subjective reports are over-reliant on the magnitude of evidence for a choice, whereas objective performance is driven by the balance of evidence between choice alternatives (Koizumi et al., 2015; Maniscalco et al., 2016; Peters et al., 2017; D. Rahnev et al., 2011; D. A. Rahnev et al., 2012; Samaha et al., 2016, 2017, 2019; Zylberberg et al., 2012). If the contents of consciousness reflect such biased computations, future work may capitalize on this dissociation to identify brain processes that down-weight unchosen alternatives. As an example of this strategy, a recent experiment recorded population activity in the superior colliculus (SC) of macaques while inducing the PEB with motion stimuli (Odegaard et al., 2018). It was found that choice-predictive activity in SC neurons was sensitive to confidence and accuracy when the two measures were correlated but was not correlated with confidence when accuracy was held constant using the PEB. Continued use of the PEB paradigm could serve as a sensitive tool to reveal neural dynamics associated specifically with subjective reports of sensory processing.

Dissociating confidence and performance has further promise in aiding research into the function of subjective perceptual states. Using the PEB in an orientation estimation task, we recently demonstrated that enhancing confidence independent of accuracy was associated with stronger serial dependencies in orientation estimation between trials (Samaha et al., 2019). This finding suggested that a candidate function of subjective confidence is enhancing temporal continuity between perceptual inputs. Using a related confidence-accuracy dissociation paradigm, Desender et al (2018) found that reducing confidence without changing performance led participants to seek out additional perceptual information before committing to a choice. This would suggest that an additional function of subjective confidence is to inform future decision-making policies. Future work will be needed to determine other functions of confidence computations as well as the ultimate perceptual or decisional locus of this bias in subjective visual report.

## Notes

Work supported by MH095984 and John Templeton Foundation grant #48365 to JS and RD

